# Effects of temperature of plant cultivation on plant palatability modify species response to novel climate

**DOI:** 10.1101/841148

**Authors:** Tomáš Dostálek, Maan Bahadur Rokaya, Zuzana Münzbergová

## Abstract

Climate warming is expected to strengthen the plant-herbivore interactions and thus increase the plant consumption rate. However, indirect impacts of temperature (acting via changes in host plant quality) on herbivore performance have only rarely been studied, and therefore, the net effect of temperature change is difficult to predict. We thus tested the effects of temperature on plant palatability and assessed whether the effects can be explained by changes in leaf traits.

We conducted multi-choice feeding experiments with six species of the genus *Impatiens* cultivated at three different temperatures in the growth chambers and in the experimental garden and also studied changes in leaf morphology and chemistry.

The leaves of *Impatiens* species were most eaten when cultivated in the temperature predicted by climate warming scenario. We found the traits related to leaf morphology (SLA, LDMC and leaf size) partly mediated the effects of temperature on leaf herbivore damage. Herbivores preferred smaller leaves with lower SLA and higher LDMC values. Results of our study suggested that elevated temperature will lead to change in leaf traits and increase their palatability. This will further enhance the levels of herbivory caused by increased herbivore pressure under climate warming.

## Introduction

Insect herbivores are one of the most important drivers of performance of plant populations ^1,2^. Under upcoming global climate change, increase in temperature is expected ^3^. The general effects of global warming on insects are relatively well documented ^4^. Higher temperatures are predicted to increase insect population densities, cause alterations in their body size, genetic composition, duration of life cycles or exploitation of host plants ^5^. Warming is thus expected to strengthen various herbivore-plant interactions ^5,6^. However, these predictions do not take into account that increased temperature might also affect the plant traits that determine their palatability to the herbivores ^7^.

Plants are growing along a wide range of biotic and abiotic conditions ^8^ and variation in herbivore damage largely depends on such conditions ^9–11^. The significant relationship between the plant damage by herbivores and the environmental conditions was shown in many previous studies. Plants tend to suffer higher herbivore damage in wetter, nutrient richer and more shaded habitats ^12–14^, i.e., in habitats in which they often grow more vigorously as suggested by Plant vigor hypothesis ^15^. Increased leaf quality also contributes to greater levels of herbivory in nutrient richer habitats ^7,16^. Leaves of plants growing in benign conditions habitats often also have higher specific leaf area, lower tissue density, thinner leaf lamina and weaker veins ^17^ and are less tough ^18^, which makes them more palatable to the herbivores. On the other hand, environmental stress can reduce plant resistance to herbivores making them more palatable according to Plant stress hypothesis ^19^. For example, ^20^ demonstrated that drought stressed plants had fewer secondary metabolites and were consequently more affected by some of their herbivores.

The effects of temperature on plant palatability are often studied along altitudinal or latitudinal gradients ^21,22^. More palatable plants at higher altitudes/latitudes suggest lower palatability at higher temperatures ^7^. The plants at lower altitudes/latitudes might be better defended ^10,23^, produce leaves morphologically harder to consume (with low specific leaf area or with trichomes ^24^) or have lower nitrogen and phosphorus leaf content ^25^. However, most of the above described studies are from natural environments where other factors except for temperature varied as well (such as intensity of UVB radiation, precipitation and abundance of herbivores). Moreover, they often do not take into account that the differences in plant traits are caused not only by different environmental condition but also by genetic differentiation of the populations ^26^. This might be the reason why not all studies find the above described relationship between altitude/latitude and plant palatability ^27–30^. Although temperature is one of the factors which is expected to change significantly under global warming, studies on its direct impact on plant palatability are still limited (but see ^7,31^).

Previous studies often found contrasting results when exploring the effects of environment on plant-herbivore interactions among different plant species indicating that plant species and environmental conditions interact to jointly determine plant palatability ^7,32–34^. To understand the effects of temperature on plant palatability, we thus need to perform studies exploring these interactive effects of plant species and temperature. By simultaneously exploring differences in plants traits, it should be possible to identify the mechanism underlying the temperature effects.

The aim of the study is to explore the effect of temperature on plant traits and palatability by insect herbivores across a range of closely related species. For the study, we use six species from genus *Impatiens*, Balsaminaceae, naturally growing in Himalayas. We selected this plant group as it is a species rich group with many species co-occurring at similar habitats along wide altitudinal ranges with strong gradient of temperature ^35^. In addition, interaction of several species from the genus *Impatiens* with their herbivores have been previously intensively studied ^36,37^ and herbivory was suggested to be one of the factors responsible for distribution of some *Impatiens* species ^38,39^. Genus *Impatiens* also contains several invasive species ^40,41^ with strong effects on native diversity ^42,43^ and bringing novel insights on drivers of plant-herbivore interactions within this group is thus highly needed.

We asked following questions: 1) What is the effect of temperature of plant cultivation on plant palatability?, 2) Can we explain the differences in plant palatability by the differences in leaf traits?, 3) What are the differences in leaf traits and palatability among the *Impatiens* species and does the effect of plant species identity interact with the effects of temperature of plant cultivation?

To answer these questions, we conducted multi-choice feeding experiments with six species of the genus *Impatiens* cultivated at three different temperatures in the growth chambers and in the experimental garden. As the herbivore, we used a generalist omnivore desert locust (*Schistocerca gregaria*) widely used in similar studies.

## Material and Methods

We used six species of genus *Impatiens*, Balsamiaceae family: *I. balsamina* L., *I. racemosa* DC., *Impatiens scullyi* Hook.f., *Impatiens tricornis* Lindl. (before revision by ^44^ usually called *I. scabrida*), *I. falcifer* Hook.f., and *I. devendrae* Pusalkar (Online Resource 1). All the species are annuals native to Himalayan region (Nepal, India) where their seeds were collected in autumn 2017. Seeds were collected from at least five mother plants in populations consisting of at least several tens of individuals. The species selection was done from a larger species collection from the region driven by leaf availability for the experiment.

Plant palatability was tested by multi-choice feeding experiments (e.g. ^45–48^). As a herbivore, we used desert locust (*Schistocerca gregaria* Forskål; Orthoptera, Acrididae) individuals of 1–2 cm, which had been purchased from a commercial insect provider (www.sarancata.cz). The desert locust is a generalist herbivore occurring in many regions of Africa, the Middle East and Asia ^49^. It is extremely polyphagous leaf chewing invertebrate and this make it excellent bioassay species for comparing leaf palatability across a wide range of plant species ^50,51^. Due to its extreme variability in food sources, it is unlikely to share a coevolutionary history with any of our study species, and their feeding preferences thus should reflect general palatability of the plants ^52^.

The experiments were done in circular arenas, 50 cm in diameter and 30 cm high. One third of the arena was filled with common garden soil. Three or six Eppendorf tubes (see below) without lids, 1.5 ml each, were regularly placed into a circle inside the arena, each tube about 10 cm from the edge. The tubes were inserted into the soil so that their top was about 1 mm above the soil surface and filled with water. One randomly chosen fully expanded leaf was placed into each Eppendorf tube with the petiole submerged in the water and the rest sticking out. Five individuals of desert locust were added into the center of the arena and the whole arena was covered with a fine mesh to prevent herbivores from escaping ^32^. For each arena, we used leaves of as similar sizes and age as possible. The experiments lasted 2-3 days and the arenas were kept under the room temperature. The duration of each experiment was based on the actual leaf damage during daily controls of the experiment. The experiment was terminated when more than 80% of at least one leaf within the arena was consumed or when more than 2/3 of the leaves had visual damage ^48^.

We performed two experiments within our study, both comparing the proportion of leaf area eaten by desert locust (leaf herbivory hereafter) of plants exposed to the herbivores in common arenas. In the first experiment (Experiment 1), each arena contained leaves of one plant species from three different temperature regimes and we used different arenas for the different plant species. In the second experiment (Experiment 2), each arena contained leaves of six different *Impatiens* species of the same origin and different arenas contained leaves cultivated under different environment. The two experiments were set up to study the effects of plant species as well as temperature of plant origin and their interaction, but at the same time to account for the technical limitations given by different phenology of the different plant species under different conditions.

### Experiment 1

In the first experiment, we primarily focused on comparing effect of cultivation in three temperature regimes in individual *Impatiens* species. *Impatiens* seeds were stratified on wet filter paper in the fridge (4°C) until germination. Triplets of germinating seeds of the same species were transplanted into 5 x 5 x 8.5 cm pots filled with a mixture of common garden soil and sand (1:2) and placed to the three growth chambers (Vötch 1014) differing in their temperature regimes. After two weeks, the seedlings were weeded to keep only one seedling per pot. There were five individuals of each of the six *Impatiens* species in each of the three growth chambers, i.e. 90 individuals in total. The temperature regimes were set to represent the present and future temperatures at localities where *Impatiens* species naturally grow in their native range in Nepal. Temperature regimes were set as follows: 1) cold regime - mean temperature from March to June in 2700 m a. s. l. representing median of higher altitudinal range of *Impatiens* species in Nepal (mean, minimum and maximum temperature of 12, 6 and 17.5°C), 2) warm regime - mean temperature from March to June in 1800 m a. s. l. representing median of lower altitudinal range of *Impatiens* species in Nepal (mean, minimum and maximum temperature of 18, 12 and 22.5°C), and 3) warm2050 regime - mean temperature from March to June in 1800 m a. s. l. representing median of lower altitudinal range of *Impatiens* species in Nepal in 2050 as predicted by global climate model MIRO5C at RCP8.5 ^53^ with mean, minimum and maximum temperature of 21, 15 and 25°C. List of *Impatiens* species in Nepal and information on their altitudinal range was obtained from Annotated Checklist of the Flowering Plants of Nepal (http://www.efloras.org/flora_page.aspx?flora_id=110), which is an updated online version of ^35^. Temperature data were obtained from Worldclim database ^54^. Data on mean temperatures in particular altitudes were obtained from slopes of correlations between altitudes and mean temperatures for particular data points along four valleys in Central and East Nepal where our seed collections took place. We used mean temperatures from March to June since it represents premonsoon period when most *Impatiens* species germinate and start to grow. For all the regimes, daily temperature course was simulated and the same day length and radiation were used, i.e. 12 h of 60% light (06.00–18.00 h; 250 μmol m^−2^ s^−1^) and 10 h of full dark with a gradual change in light availability in the transition between the light and dark period over 1 h. Pots were regularly watered with tap water. In each arena (see above), there were three leaves of the same *Impatiens* species with each of them originating from different temperature regime (growth chamber). Each plant species was studied in 10 arenas (10 replicates). Leaves were sampled in June 2018 when the plants were already grown up and flowering.

### Experiment 2

For the second experiment, we primarily focused on comparing the palatability of six *Impatiens* species cultivated under different regimes. We used leaves from the cold regime used above and from common garden environment where the plants had more natural conditions (natural light and temperature conditions, more space). Experimental garden (49°59’38’’ N, 14°33’57’’ E) is located 320 m above sea level in the temperate climate zone, with a mean annual temperature of 8.6°C and precipitation of 610 mm. Mean, minimum and maximum temperature in August (one month before leaf collection) were 20.7, 7.8 and 35.5°C, respectively. Seeds at the common garden environment were sown into 5L pots with the same soil as for Experiment 1 in January 2018 and thus they were naturally stratified. In April the seedlings were weeded to keep only one seedling per pot. There were five plants of each of the six species in each of two environments, i.e. 60 plants in total. Pots were regularly watered. In the cold regime in the growth chamber, leaves were collected for arena tests in June as in Experiment 1. In the common garden, leaves were collected in September when all the plants were grown up and flowering. In each arena (see above), there were six leaves from different *Impatiens* species. We used 10 arenas for each environment, i.e. 20 replicates (arenas) in total.

### Leaf traits

The leaves were individually fresh weighted and scanned both before and after the herbivory (e.g. ^48^). Leaf area was estimated using ImageJ software (version 1.52a, Java 1.8.0_112, Wayen Rasband, U.S. National Institutes of Health, Bethesda, MD, USA; website: http://rsb.info.nih.gov/ij/download.html). After herbivory, the leaves were dried to a constant weight and weighted again. This information, together with information on leaf size, was used to calculate specific leaf area (SLA; mm^2^ mg^−1^ dry mass) and leaf dry matter content (LDMC; g dry mass g^−1^ fresh mass) for each leaf. Leaves were evenly eaten by herbivores and no leaf parts were preferred. In two cases the whole leaf was eaten. Then we used mean SLA and LDMC values of the other nine leaves from respective temperature regime. We also analyzed total carbon (C), nitrogen (N) and phosphorus (P) content in leaf biomass. Since it was not possible to analyze leaf nutrient content in the leaves used directly in multi-choice feeding experiments due their small size, we used mixed sample of ten randomly chosen leaves per each species and growth chamber temperature regime/common garden which were not exposed to herbivores. The chemical analyses were performed in the Analytical laboratory of the Institute of Botany, the Czech Academy of Sciences, Průhonice. The contents of nitrogen and carbon were analyzed following ^55^. The content of phosphorus was analyzed spectrophotometrically following ^56^. These traits were chosen because they encompass a range of mechanical and chemical properties that are quantifiable within and between plant species and are related to plant palatability ^57^.

### Data analyses

Data from the two experiments were analyzed separately. Leaf herbivory was square root transformed, SLA, LDMC and initial leaf area were log-transformed to meet test assumptions.

### Experiment 1

First, we tested the effect of temperature regime (growth chamber), species identity and their interaction on leaf herbivory. We used linear mixed effect model with arena code as a random factor for this test. Temperature regime was used as a factor in this and all the following tests since the effects of temperature were not linear.

Second, we tested how the leaf traits (SLA, LDMC, initial leaf area) are affected by the temperature regime, the *Impatiens* species identity and their interaction using ANOVA. Differences in leaf nutrient content were not tested since these data were not replicated within growth chambers and species variant due to lack of the plant material available for analyses. All the traits were, however, used to explain leaf herbivory (see below). Since leaf nutrient contents and their ratios were largely correlated, we only used those which were not highly correlated with each other (r < 0.9). We thus only used content of C and ratios of C:N, N:P and C:P.

Further, we explored the effect of leaf traits on leaf herbivory, either alone or after accounting for species identity and temperature regime. For these tests, we added combination of species identity and temperature regime as another random factor to account for the fact that the traits measured within one species and temperature regime are not independent and in case of leaf nutrient content they were only measured once. Usefulness of adding the traits into the models was assessed using AIC criteria leading to identification of optimal model.

### Experiment 2

The tests largely followed the logic described above in Experiment 1. First, we tested the effect of the species identity, environment and their interaction on leaf herbivory. We used linear mixed effects model with arena code as a random factor for this test. Second, we tested how the leaf traits differed among the six *Impatiens* species and the two environments (growth chamber and common garden) using ANOVA. We also explored the effect of leaf traits on leaf herbivory, either alone or after accounting for species identity and environment. Optimal model explaining leaf herbivory was constructed in the same way as in Experiment 1.

## Results

### Effect of temperature and species identity on leaf herbivory

Results of Experiment 1 showed that leaf herbivory differed among the three temperature regimes (P = 0.003, Fig. 1A) and among the six *Impatiens* species (P < 0.001, Online Resource 2). Moreover, there was a significant interaction between species and temperature regime (P = 0.007, Online Resource 4A). Leaves of five species were the most eaten in the warm2050 regime and most of them the least eaten in the cold one. The only exception was *I. balsamina* with the highest herbivore damage when taken from the cold temperature regime (Online Resource 4A).

**Fig. 1.**
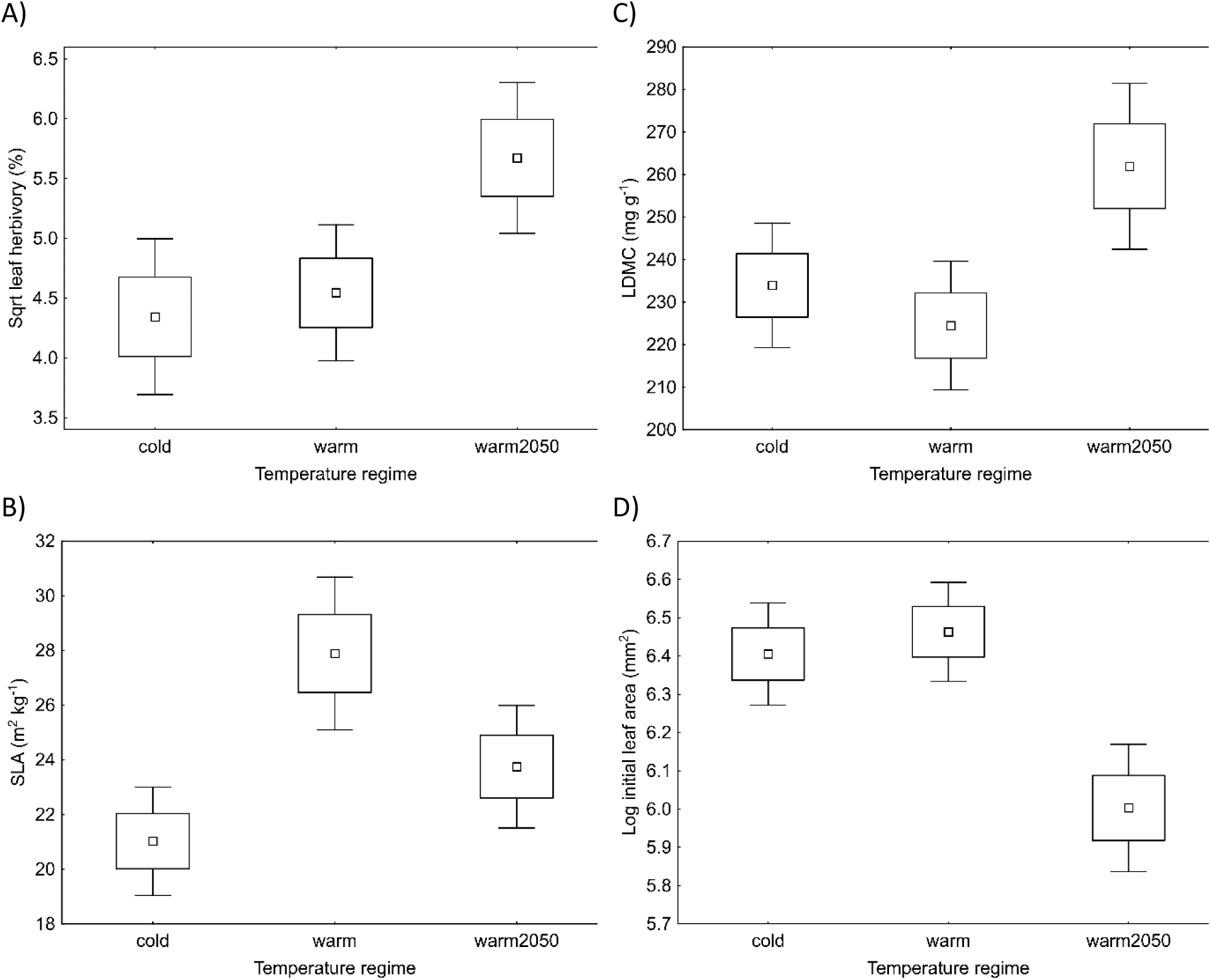
Effect of temperature regime on A) leaf herbivory, B) SLA, C) LDMC, and D) initial leaf area in Experiment 1. Box plots show means, SE and the whiskers indicate 1.96*SE.

In Experiment 2, leaf herbivory also differed among the six *Impatiens* species (P = 0.022) but was not affected by the environment (P = 0.194) or the interaction between species identity and environment (P = 0.236). *I. balsamina* and *I. racemosa* tended to be the least damaged by the herbivores (Fig. 2, Online Resource 3).

**Fig. 2.**
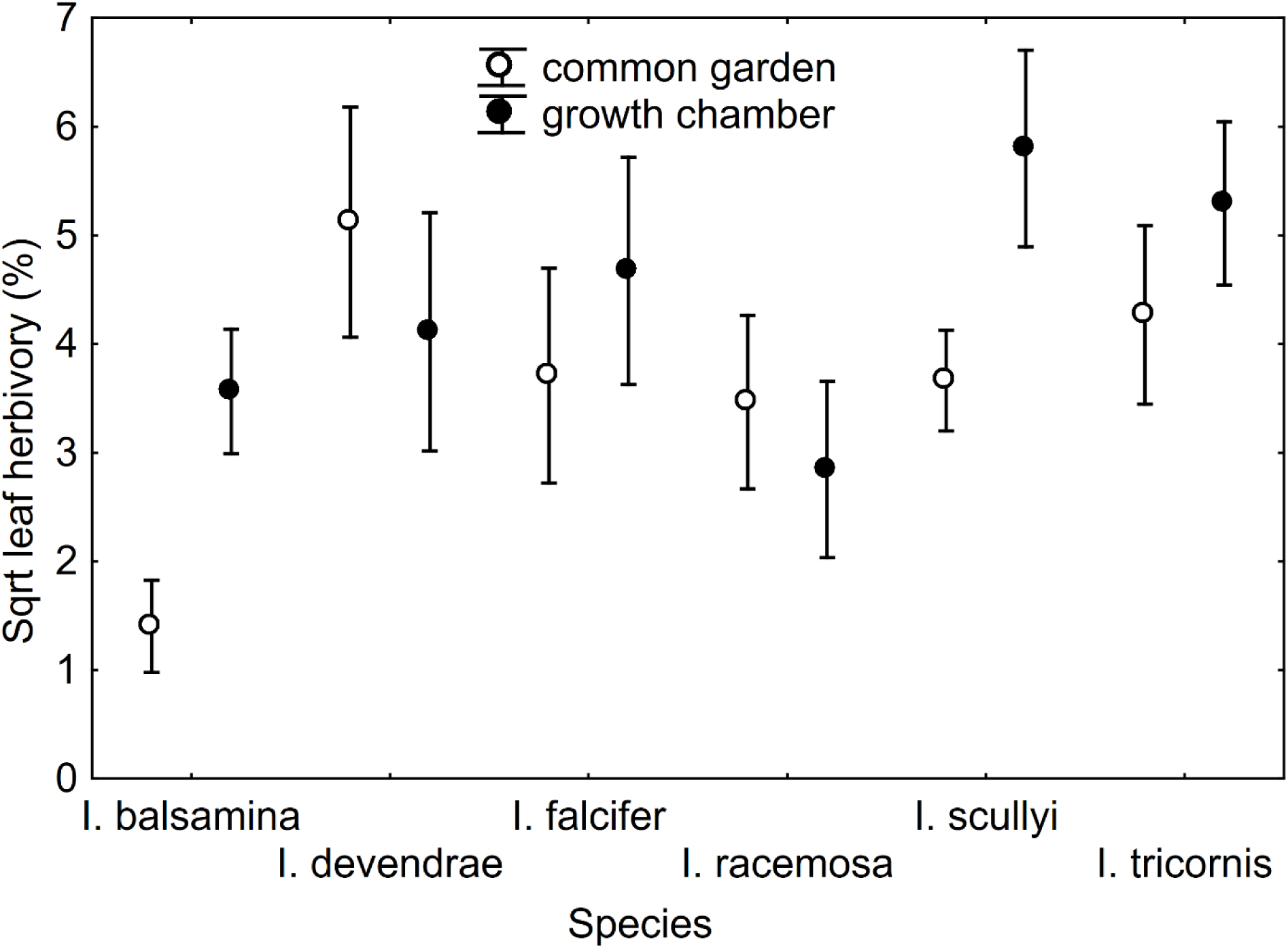
Differences in leaf herbivory among six *Impatiens* species between the two environments (common garden vs. growth chamber) in Experiment 2. Means and their standard errors are shown.

### Effect of temperature and species identity on leaf traits

In experiment 1, all the tested leaf traits significantly differed among the temperature regimes and among species (P < 0.002 in all cases) (Online Resource 2). The highest values of SLA were recorded in the warm and the highest LDMC in the warm2050 temperature regime (Fig. 1B-C). Initial leaf area was similar in the cold and warm regime and was the lowest in the warm2050 regime (Fig. 1D). However, significant interaction between the temperature regime and species identity in all tested traits indicated different trends among the six species (Online Resource 2, Online Resource 4B-D). Leaf nutrient contents (C, C:N, C:P, N:P) did not show consistent trend among temperature regimes across the studied species (Online Resource 2, Online Resource 3).

In Experiment 2, the six *Impatiens* species differed significantly in SLA, LDMC and initial leaf area (P < 0.005 in all cases) and there were also significant differences between the two environments, i.e., the growth chamber and common garden (P < 0.001 in all cases, Online Resource 4B-D). Similarly to the first experiment, significant effects of interactions between environment and species identity indicated different trends in leaf sizes (P < 0.001) and SLA (P = 0.048) among *Impatiens* species between common garden and growth chamber. Interaction between LDMC and environment was not significant (P = 0.66, Online Resource 3).

### Importance of leaf traits for leaf palatability

In Experiment 1, SLA, LDMC and the initial leaf area significantly contributed to explaining leaf herbivory compared to the null model including only the effects of temperature regime and species identity. The optimal model explaining leaf herbivory included SLA, LDMC and initial leaf area (Table 1A). Herbivores preferred smaller leaves but their response to SLA and LDMC was not consistent across the six *Impatiens* species (Fig. 3). When the null model did not include temperature regime and species identity, the optimal model included LDMC, initial leaf area and C:N ratio. SLA and the other leaf nutrient contents were not included in the optimal models explaining variation in leaf herbivory (Table 1A).

**Table 1.**
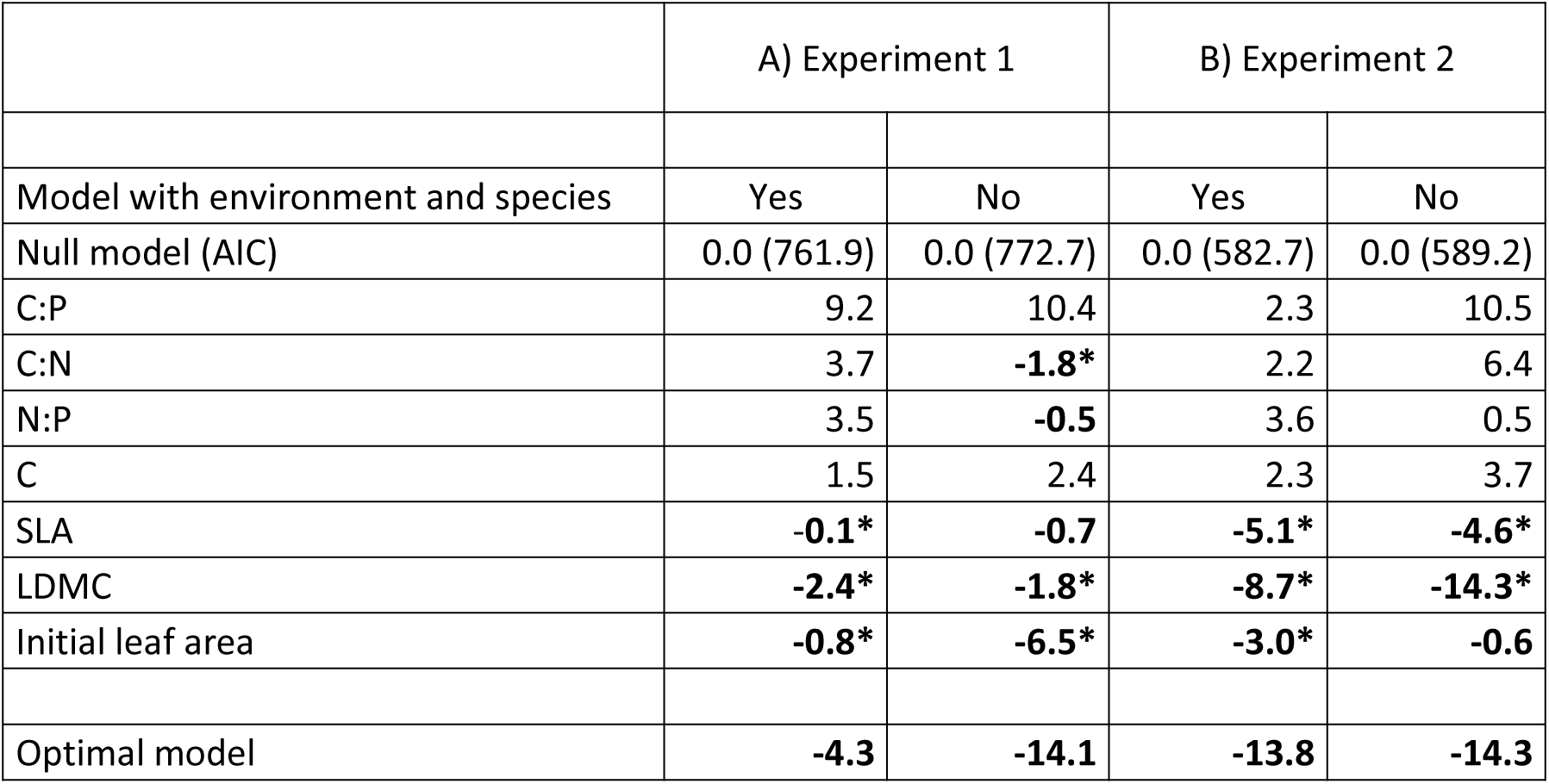
Leaf traits explaining leaf herbivory with or without accounting for the effect of environment and species in the null model. Environment is represented A) by three different temperature regimes in Experiment 1 and B) by common garden vs. growth chamber in Experiment 2. The values shown are values of ΔAIC comparing the model with the given trait to the null model. Numbers in the brackets show AIC values of the null model. Leaf traits improving the model (decreasing model AIC) are in bold. Arena and combination of environment and species were used as random factors in all the tests. Asterisks indicate leaf traits included in the optimal model.

**Fig. 3.**
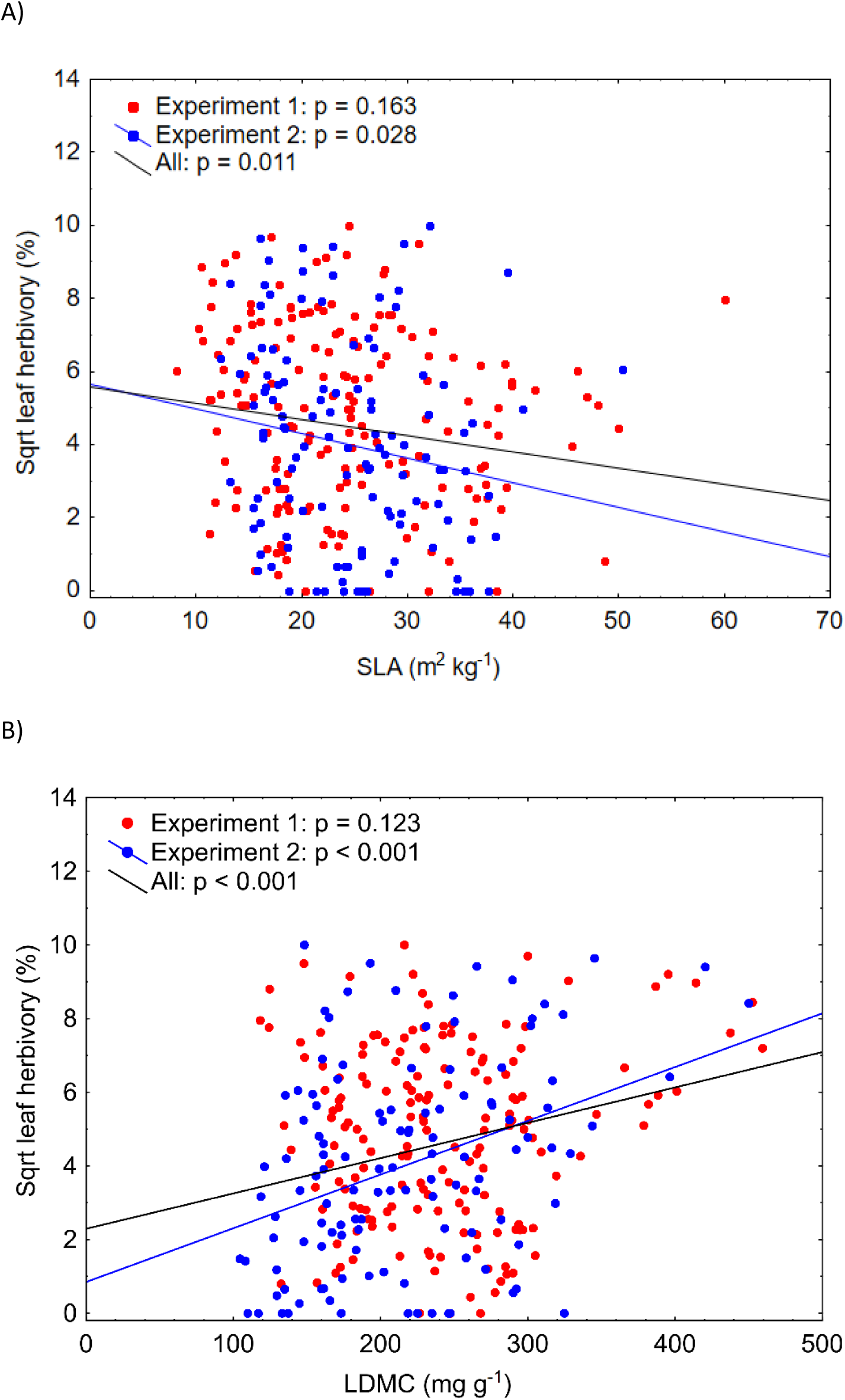
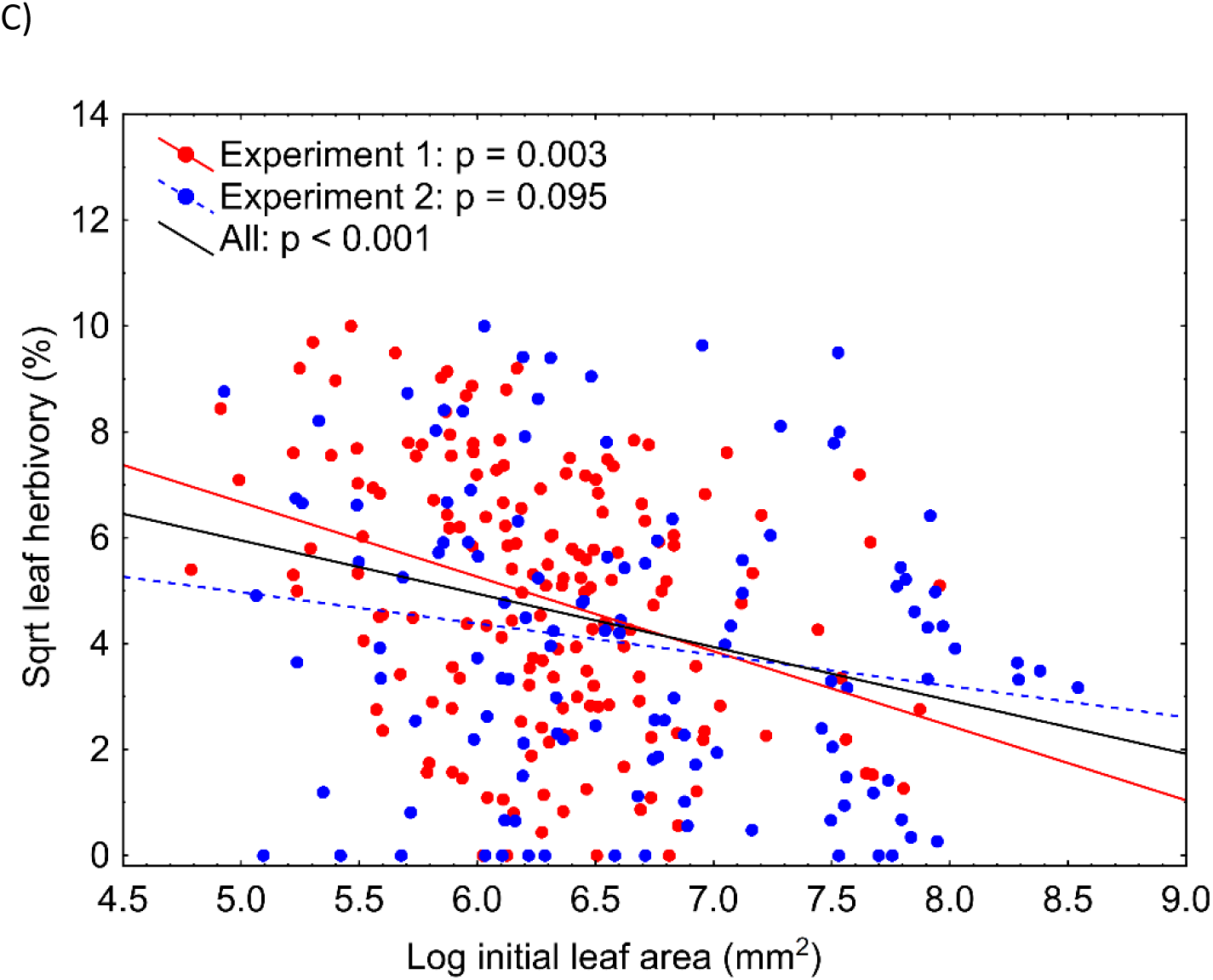
Relationship between leaf herbivory and A) SLA, B) LDMC and C) initial leaf area using data from Experiment 1 (in red) and Experiment 2 (in blue). Data on individual leaves are presented. Solid and dashed lines show significant (P < 0.05) and marginally significant (P < 0.1) relationship, respectively. Fitted line corresponds with the tests for Experiment 1 and 2 presented in Table 1 without including the effect of environment and species. Black line represents fit for all the data combined.

Similarly to Experiment 1, SLA, LDMC and initial leaf area significantly contributed to explaining leaf herbivory compared to the null model including only species and environment in Experiment 2. The optimal model explaining leaf herbivory included all these three traits (Table 1B). Herbivores preferred smaller leaves and leaves with smaller SLA and higher LDMC (Fig. 3). When null model did not include species and environment, the optimal model only included SLA and LDMC. Leaf nutrient contents did not explain any additional variation in leaf herbivory in both models in Experiment 2 (Table 1B).

## Discussion

Global warming is expected to strengthen the plant-herbivore interactions and thus increase the plant consumption rate especially by ectotherm omnivores ^27,58^. However, indirect impacts of temperature (acting via changes in host plant quality) on herbivore performance have only rarely been studied, and therefore, the net effect is difficult to predict. Our study is one of the very few exploring both effects of temperature of plant cultivation and leaf traits on herbivore damage. We found strong effect of temperature on herbivore damage which can be partly explained by temperature effects on leaf traits mainly related with leaf morphology such as SLA, LDMC and leaf size. We also showed that the effect of elevated temperature on plant palatability strongly differ among the *Impatiens* species and conclusions about the effects of climate warming must be done specifically for each species.

### Effect of temperature on leaf herbivory

Results of our study suggest that plant palatability increases with temperature. This is in contrast with previous study of ^7^. They found that rising temperatures (15, 20 and 25°C) significantly decreased plant palatability in aquatic plant *Potamogeton lucens*, which could be explained by changes in the underlying leaf nutrient content (decreasing content of N and P in the leaves). However, they did not find any effect of temperature on palatability in two other aquatic plants included in their study. The absence of an apparent relationship between temperature and leaf nutrient content in our study could explain contrasting pattern. Other studies on the effect of elevated temperature on herbivore damage did not find any significant result at *Salix myrsinifolia* ^59^ and *Quercus pubescens* ^31^. They admitted that the reason might be very small temperature increase (only about 2°C) which was not enough to affect leaf traits affecting species palatability.

### Effect of temperature on leaf traits

We found that temperature regime affected all measured leaf traits. Many studies reported positive relationship between temperature and SLA (e.g., ^60–62^. It could be attributed to plants’ ability to change leaf thickness or cell size depending on environmental conditions ^63^. In our study, however, it only happened when temperature increased between cold and warm growth chamber. In the growth chamber with elevated temperature according to the climate warming scenario (warm2050), SLA decreased again. Even though many studies predict linear relationship between SLA and temperature (e.g., ^62,64^, studies looking on response of individual species often show variable results. ^60^ report non-linear responses in SLA to temperature in several species similar to the pattern detected in our study. It may be because the highest temperature is out of the species optima leading to changes in leaf structure. LDMC showed similar but inversed pattern to SLA, which is in agreement with other studies ^62,64,65^. We also found that leaf size decreased with increasing temperature, especially in warm2050 regime. This might be at least partly related to non-linear relationship between SLA/LDMC and temperature. Larger leaves are not advantageous in elevated temperatures as they increase transpiration area ^66^. Surprisingly, we did not find any consistent effect of temperature on leaf nutrient content as for example ^25^, ^67^ and ^7^ who found lower nitrogen and phosphorus leaf content in plants growing at higher temperatures. *Impatiens* seems to respond to elevated temperature by change in leaf morphological structures while keeping the nutrient content in the leaves at the similar level. However, more data are needed to confirm this as our leaf nutrient content data were not replicated and thus could not be formally tested. Similarly to our study, meta-analysis of ^27^ showed no effects of temperature increase on either nitrogen concentrations or C/N ratio and they suggested that not nutrients but defense chemicals are the main reason for lower palatability of plants at elevated temperatures.

### Importance of leaf traits for leaf palatability

We demonstrated that herbivores prefer plants with lower SLA and higher LDMC. This contrasts with the conclusions of several previous studies demonstrating strong positive effect of SLA on leaf herbivory e.g. ^34,68,69^. High SLA is primarily related to growth rate and resource acquisition, but it is also expected to contribute to greater palatability to herbivores ^68,70^. The difference in the results may be caused by the fact that the previous studies used wider ranges of species with more variable SLA as suggested by ^32^ who also found negative correlation between herbivore damage and SLA. They further argued that the negative correlation might be due to leaf sampling throughout range of different habitats which was not the case in our study when all the plants were grown in the same conditions only differing in temperature. Very high SLA values at *Impatiens* species might be another reason for non-positive relationship between SLA and herbivory. Compared to other studies (such as ^32,34,68^), gradient of SLA started at higher values (indicating thin leaves) and the highest SLA values recorded at our study (50 m^2^kg^−1^) probably are not attractive for herbivores any more due to being too thin. Thus, the direction of the relationship between palatability and SLA may in fact be unimodal on large scale and the direction detected in the different studies depends on the exact range of the SLA values.

Our results also indicate that herbivores preferred smaller leaves. This is in contrast with many other studies which found that more vigorous plants suffered more leaf damage e. g. ^71,72^ as predicted by Plant Vigour Hypothesis ^15^. Similarly to our study, ^73^ found higher herbivory of gall forming insect at smaller and not larger leaves. They suggested that smaller leaves should possess higher concentrations of resources essential for larval development. However, we found higher nitrogen and phosphorus content (and less carbon) in leaf biomass of larger leaves even though these relationships are based on means at the population and environment level (Fig. S3). Negative relationship between leaf size and extent of herbivore damage might be due to increase in concentration of substances decreasing leaf palatability at larger leaves as found by ^74^. They found that decrease in palatability at larger willow seedlings was positively correlated with an increase in condensed tannin concentration. ^75^ also suggested that young (i.e. small) leaves of *Phytolacca americana* may be more nutritious and less tough than mature leaves explaining greater herbivory.

Even though ratios C:N and N:P were included in the optimal model explaining leaf herbivory, they did not explain any additional information when temperature regime and species identity were included in the model indicating that their effects cannot be distinguished from the effects of temperature and species. In agreement, review of ^76^ tested the importance of plant traits for plant-herbivore interactions and they concluded that morphological and physical traits are often more important for plant-herbivore interactions than chemical traits.

### Differences in response to elevated temperature among Impatiens species

Most *Impatiens* species in our study responded similarly to elevated temperature. The only exception was *I. balsamina*. In contrast to all the other species, *I. balsamina* increased SLA and decreased LDMC under the warmest (warm2050) temperature regime (Online Resource 4). Moreover, leaves of all other species were most eaten when cultivated in the warmest temperature regime, while leaves of *I. balsamina* were most eaten when cultivated in the cold temperature regime. *I. balsamina* was also the least eaten species overall. However, this effect was only significant when the plants were cultivated in the common garden environment with higher temperatures comparable with the warmest regime in the growth chambers. When we compared the species cultivated in cold temperature regime in the growth chamber, there were no differences in their palatability. The reason for the strong differences in the warmest regime might be that original localities of all species included in our study are at around 2500 m a. s. l. and warm2050 regime represents temperature under predicted climate change for 1800 m a. s. l. (median of lower altitudinal range of all *Impatiens* species in Nepal – see methods for details). The only exception is *I. balsamina*, seeds of which were collected at 1330 m a. s. l., which is altitude with temperatures simulated in warm2050 regime in our study. *I. balsamina* thus seems to be well adapted to temperatures at its home localities. Compared to other species, its leaves are tougher and might be better adapted to retain water under elevated temperature. Compared to other species, leaves of *I. balsamina* are also less palatable to herbivores under elevated temperature. ^77^ recorded increased tannin concentration in leaves of *I. balsamina* following drought stress and suggested they might play a role in its resistance against insect herbivores. While the other *Impatiens* species included in our study are expected to suffer from higher herbivory under elevated temperature, invasive potential of *I. balsamina* is larger under predicted global climate warming as suggested also by ^78^. The results thus indicate that changes in palatability due to changes in temperature may contribute to reduced performance in cold-adapted species under future climate warming, while at the same time it may increase the success of warm-adapted species.

## Acknowledgements

The study was supported by the Czech Science Foundation (project no. 17-10280S) and long-term research development project no. RVO 67985939. Participants in the POPEKOL seminars and Hana Skálová provided us with many useful comments and Wojciech Adamowski helped us with species identification. We are thankful to Ilona Jarošincová, Zuzana Líblová and Martina Lokvencová for helping us during the plant cultivations and multi-choice feeding experiments.

## Author Contributions

TD and ZM conceived and designed the experiments. MBR collected the plant material in the field. TD performed the experiments. TD and ZM analyzed the data. TD and ZM wrote the manuscript.

## Data Availability

The data reported in this paper has been deposited at FigShare (https://doi.org/10.6084/m9.figshare.9438722.v1).

