## Supplemental information for "Effects of temperature of plant cultivation on plant palatability modify species response to novel climate"

**Online Resource 1** Description of six *Impatiens* species used in the study. Locality name, GPS coordinates (WGS 84) and altitude are shown for localities where the seeds were collected in autumn 2017. Mean premonsoon temperatures were obtained from Worldclim database (Hijmans et al. 2005) as mean temperatures from March to June for particular localities in 1960-90. Premonsoon period represents time when most *Impatiens* species germinate and start to grow. Information on plant height, species altitudinal range and habitat types are species characteristics from the literature (Akiyama et al. 1991; Grey-Wilson 1991; Pusalkar and Singh 2010; Akiyama and Ohba 2016).

| Species | Locality name | GPS coordinates | | Altitude (m a.s.l.) | Mean premonsoon temperature (°C) | Plant height (cm) | Species altitudinal range (m a.s.l.) | Habitat type |
|---|---|---|---|---|---|---|---|---|
| | | N | E | | | | | |
| *I. balsamina* L. | Kirtipur, Kathmandu | 27.68377 | 85.28356 | 1330 | 20.4 | 60 | 200-1800 | Widely naturalized and cultivated in Himalayan foothills. |
| *I. devendrae* Pusalkar* | Darchula | 29.89019 | 80.92719 | 2728 | 10.9 | 30-80 | 2400-3200 | Shaded or partly shaded, moist places in Rhododendron forests and along forest edges. |
| *I. falcifer* Hook.f. | Phulchowki, Lalitpur | 27.57744 | 85.39950 | 2499 | 15.2 | 10-50 | 2300-3200 | Montane forests and stream beds. |
| *I. racemosa* DC. | Chandragiri | 27.66517 | 85.20458 | 2525 | 14.2 | 10-60 | 1300-3900 | Broad-leaved forest, rocky places, path sides, stream margins. |
| *Impatiens scullyi* Hook.f. | Chame, Manang | 28.55227 | 84.24152 | 2688 | 12.8 | 30-60 | 2000-3600 | Forest understories, thickets along riverbanks, shaded moist places. |
| *Impatiens tricornis* Lindl. ** | Chame, Manang | 28.55227 | 84.24152 | 2688 | 12.8 | 30-80 | 1000-3600 | Forest understories, thickets along riverbanks, shaded moist places. |

* Proper species determination is still in progress. Herbarium specimen was collected, and species identity is currently being clarified using molecular data.

** Before revision by (Akiyama and Ohba 2016) usually called *I. scabrida.*


Akiyama S, Ohba H (2016) Studies of Impatiens (Balsaminaceae) of Nepal 3. Impatiens scabrida and Allied Species. Bull Natl Mus Nat Sci Ser B Bot 42:121–130

Akiyama S, Ohba H, Wakabayashi M (1991) Taxonomic notes of the East Himalayan species of Impatiens. Studies of Himalayan Impatiens (Balsaminaceae). In: Ohba H, Malla SM (eds) The Himalayan Plants 2. University of Tokyo Press, Tokyo, pp 66–94

Grey-Wilson C (1991) Balsaminaceae. In: Grierson AJC, Long DG (eds) Flora of Bhutan. Royal Botanic Garden, Edinburgh, UK, pp 82–104

Hijmans RJ, Cameron SE, Parra JL, et al (2005) Very high resolution interpolated climate surfaces for global land areas. Int J Climatol 25:1965–1978. doi: 10.1002/joc.1276

Pusalkar PK, Singh DK (2010) Three New Species of Impatiens (Balsaminaceae) from Western Himalaya, India. 55:11

**Online Resource 2** Leaf herbivory, leaf traits and leaf nutrient contents in three temperature regimes in six *Impatiens* species in Experiment 1. The same letters in the *Sig* columns indicate non-significant differences among temperature regimes within species (P > 0.05). Differences in leaf nutrient content were not tested since these data were not replicated within growth chambers and species variant due to lack of the plant material available for analyses.

| Species | Temp regime | Leaf herb. (%) | Sig | Initial leaf area ($mm^2$) | Sig | SLA $m^2 kg^{-1}$ | Sig | LDMC $mg g^{-1}$ | Sig | N (%) | C (%) | P (%) | C:N | C:P | N:P |
|---|---|---|---|---|---|---|---|---|---|---|---|---|---|---|---|
| *I. balsamina* | cold | 50.0 | a | 616.3 | b | 0.17 | ab | 19.22 | a | 1.77 | 37.18 | 0.18 | 21.05 | 201.81 | 9.59 |
| | warm | 34.3 | a | 486.3 | b | 0.13 | a | 28.86 | b | 1.24 | 37.65 | 0.16 | 30.31 | 237.74 | 7.84 |
| | warm2050 | 43.1 | a | 325.4 | a | 0.21 | b | 20.71 | a | 1.65 | 38.30 | 0.19 | 23.24 | 197.10 | 8.48 |
| *I. racemosa* | cold | 15.4 | a | 616.1 | a | 0.35 | a | 18.34 | a | 1.16 | 39.09 | 0.16 | 33.84 | 249.81 | 7.38 |
| | warm | 28.9 | b | 639.2 | a | 0.40 | a | 18.23 | a | 1.03 | 36.00 | 0.15 | 34.95 | 239.18 | 6.84 |
| | warm2050 | 33.8 | b | 505.0 | a | 0.36 | a | 21.09 | a | 1.75 | 41.08 | 0.12 | 23.42 | 339.87 | 14.51 |
| *I. scullyi* | cold | 9.0 | a | 1471.6 | a | 0.20 | ab | 29.04 | a | 1.16 | 36.96 | 0.19 | 31.91 | 199.05 | 6.24 |
| | warm | 14.1 | a | 1470.9 | a | 0.24 | b | 29.01 | a | 1.65 | 36.67 | 0.19 | 22.16 | 193.13 | 8.71 |
| | warm2050 | 42.2 | b | 1323.4 | a | 0.18 | a | 37.37 | b | 1.73 | 37.86 | 0.19 | 21.83 | 199.10 | 9.12 |
| *I. tricornis* | cold | 28.4 | a | 534.8 | ab | 0.18 | a | 23.14 | a | 1.32 | 36.63 | 0.20 | 27.84 | 186.89 | 6.71 |
| | warm | 25.2 | a | 773.6 | b | 0.29 | c | 20.65 | a | 1.97 | 37.54 | 0.27 | 19.02 | 139.72 | 7.35 |
| | warm2050 | 39.4 | a | 385.9 | a | 0.22 | b | 24.72 | a | 1.40 | 38.02 | 0.12 | 27.14 | 327.28 | 12.06 |
| *I. falcifer* | cold | 39.1 | ab | 453.3 | b | 0.18 | a | 24.83 | b | 1.51 | 37.38 | 0.20 | 24.72 | 185.10 | 7.49 |
| | warm | 25.2 | a | 428.1 | b | 0.31 | b | 19.01 | a | 1.06 | 37.51 | 0.14 | 35.36 | 261.19 | 7.39 |
| | warm2050 | 49.0 | b | 188.9 | a | 0.17 | a | 31.50 | c | 1.24 | 37.34 | 0.13 | 30.16 | 290.38 | 9.63 |
| *I. devendrae* | cold | 3.6 | a | 738.2 | ab | 0.18 | a | 27.50 | b | 0.79 | 37.85 | 0.11 | 48.13 | 345.03 | 7.17 |
| | warm | 21.0 | b | 822.1 | b | 0.30 | b | 20.91 | a | 1.02 | 37.71 | 0.23 | 36.86 | 166.40 | 4.51 |
| | warm2050 | 21.6 | b | 592.5 | a | 0.27 | b | 25.13 | b | 1.19 | 38.71 | 0.17 | 32.48 | 224.06 | 6.90 |

**Online Resource 3** Leaf herbivory, leaf traits and leaf nutrient contents in six *Impatiens* species in common garden and growth chamber environment in Experiment 2. The same letters indicate non-significant among the species within environment (P > 0.05). Differences in leaf nutrient content were not tested since these data were not replicated within growth chambers and species variant due to lack of the plant material available for analyses.

| Species | Temp regime | Leaf herb. (%) | Sig | Initial leaf area ($mm^2$) | Sig | SLA $m^2$ $kg^{-1}$ | Sig | LDMC mg $g^{-1}$ | Sig | N (%) | C (%) | P (%) | C:N | C:P | N:P |
|---|---|---|---|---|---|---|---|---|---|---|---|---|---|---|---|
| *I. balsamina* | Garden | 3.6 | a | 1944.2 | d | 0.29 | a | 12.10 | a | 2.47 | 33.81 | 0.30 | 13.69 | 114.25 | 8.34 |
| *I. racemosa* | | 17.7 | ab | 700.2 | b | 0.29 | a | 16.82 | b | 2.41 | 41.18 | 0.26 | 17.09 | 155.48 | 9.10 |
| *I. scullyi* | | 15.4 | ab | 3282.6 | e | 0.30 | a | 24.03 | c | 2.39 | 36.72 | 0.25 | 15.39 | 146.95 | 9.55 |
| *I. tricornis* | | 24.3 | ab | 2296.5 | de | 0.27 | a | 19.47 | b | 2.66 | 33.94 | 0.30 | 12.77 | 112.19 | 8.79 |
| *I. falcifer* | | 22.5 | ab | 455.1 | a | 0.32 | a | 14.81 | b | 2.99 | 34.67 | 0.27 | 11.59 | 127.59 | 11.01 |
| *I. devendrae* | | 36.4 | b | 1345.5 | c | 0.27 | a | 21.64 | c | 2.32 | 34.84 | 0.33 | 14.99 | 106.06 | 7.08 |
| *I. balsamina* | Growth chamber | 15.7 | a | 825.9 | d | 0.16 | a | 17.00 | a | 1.77 | 37.18 | 0.18 | 21.05 | 201.81 | 9.59 |
| *I. racemosa* | | 14.0 | a | 518.8 | bc | 0.21 | bc | 24.50 | bc | 1.16 | 39.09 | 0.16 | 33.84 | 249.81 | 7.38 |
| *I. scullyi* | | 41.0 | a | 847.6 | cd | 0.19 | c | 31.55 | d | 1.16 | 36.96 | 0.19 | 31.91 | 199.05 | 6.24 |
| *I. tricornis* | | 33.1 | a | 341.9 | b | 0.23 | c | 24.37 | bd | 1.32 | 36.63 | 0.20 | 27.84 | 186.89 | 6.71 |
| *I. falcifer* | | 31.7 | a | 191.0 | a | 0.22 | c | 22.16 | ab | 1.51 | 37.38 | 0.20 | 24.72 | 185.10 | 7.49 |
| *I. devendrae* | | 27.7 | a | 568.4 | bd | 0.17 | b | 30.37 | cd | 0.79 | 37.85 | 0.11 | 48.13 | 345.03 | 7.17 |

**Online Resource 4** Effect of temperature regime on A) leaf herbivory, B) SLA, C) LDMC, and D) initial leaf area in multichoice feeding experiment categorized by the six *Impatiens* species in Experiment 1. Means and their standard errors are shown. *** P < 0.001, ** P < 0.01, * P < 0.05, . P < 0.1

A)

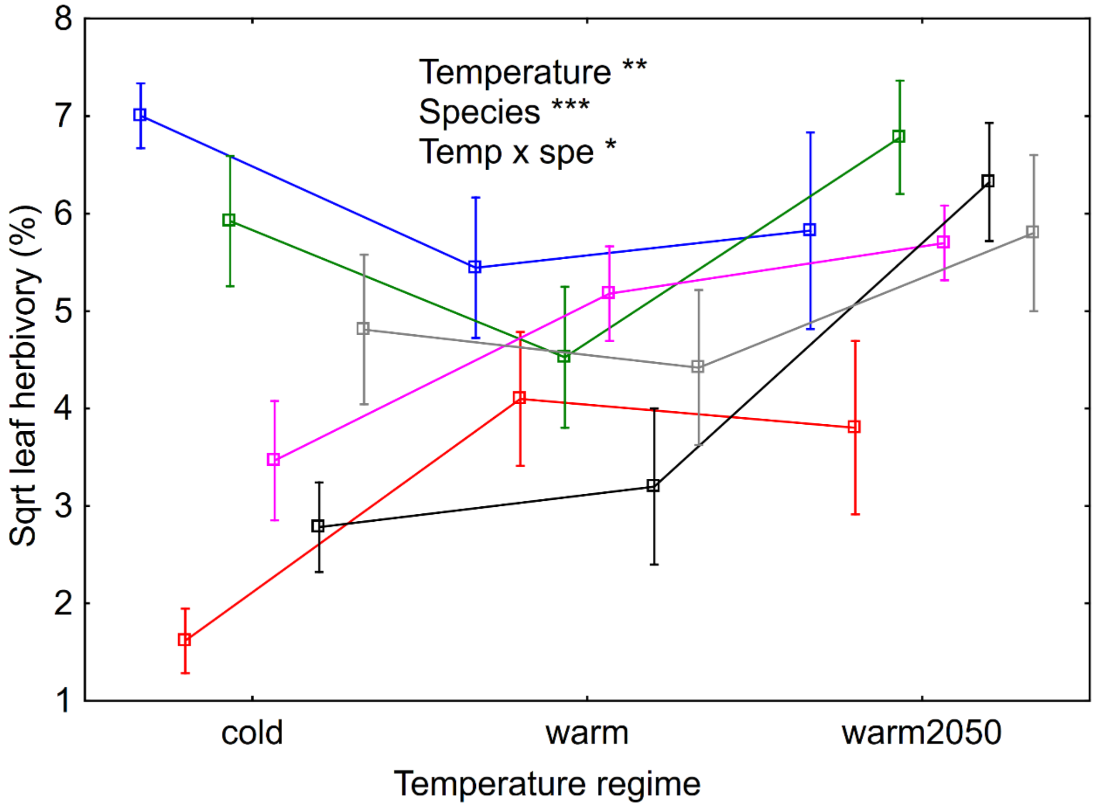

B)

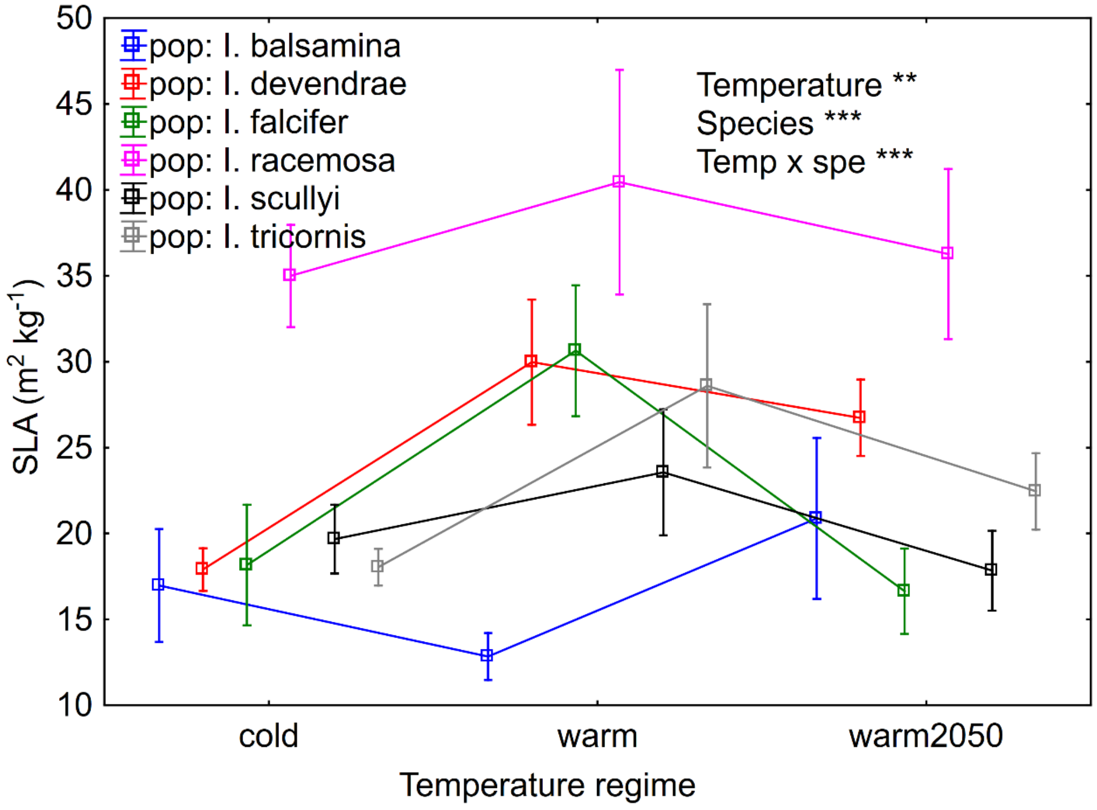

pop: I. balsamina
pop: I. devendrae
pop: I. falcifer
pop: I. racemosa
pop: I. scullyi
pop: I. tricornis
Temperature **
Species ***
Temp x spe ***
SLA (m2 kg-1)
50
45
40
35
30
25
20
15
10
cold
warm
warm2050
Temperature regime


C)

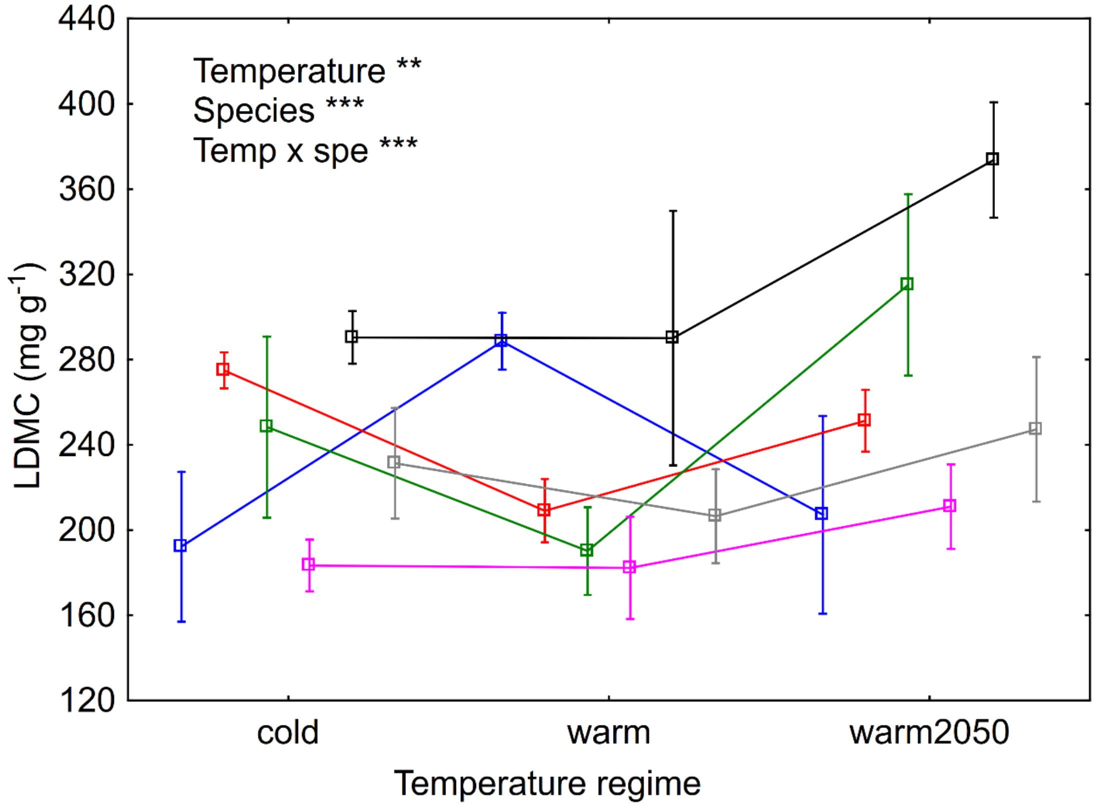

Temperature **
Species ***
Temp x spe ***
LDMC (mg g-1)
440
400
360
320
280
240
200
160
120
cold
warm
warm2050
Temperature regime

D)

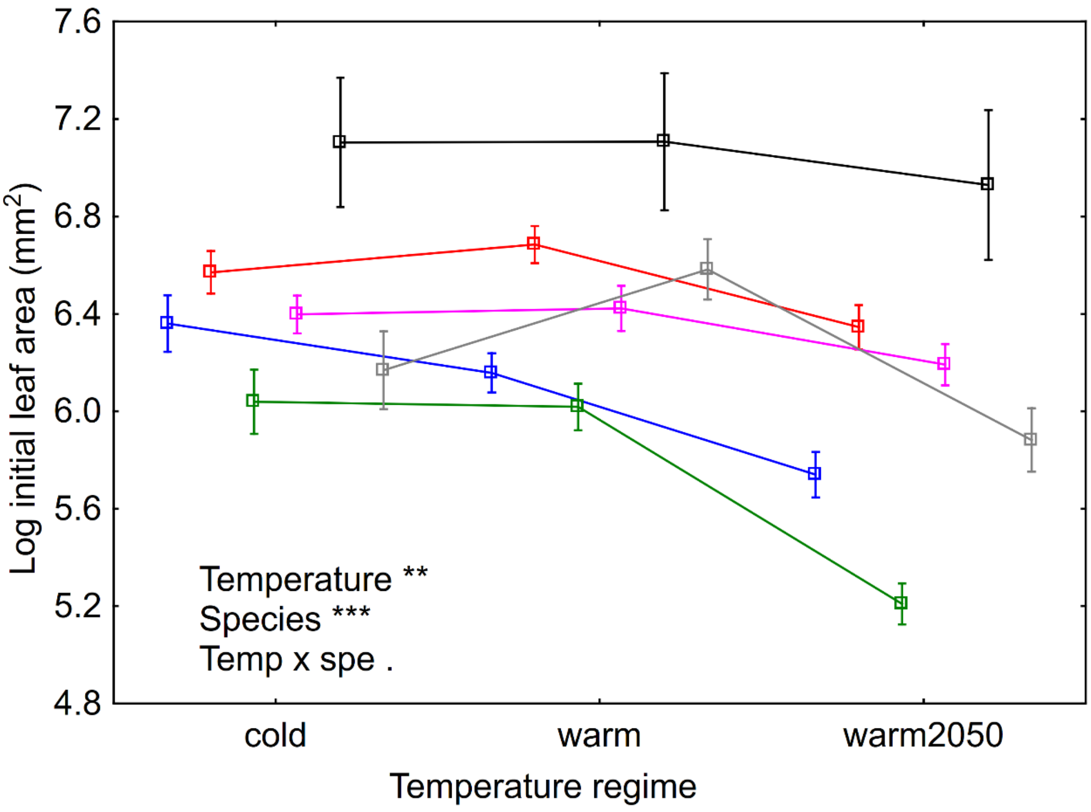

7.6
7.2
6.8
6.4
6.0
5.6
5.2
4.8
Log initial leaf area (mm2)
Temperature **
Species ***
Temp x spe .
cold
warm
warm2050
Temperature regime

**Online Resource 5** Differences in A) SLA, B) LDMC, and C) initial leaf area among six *Impatiens* species between the two environments (common garden vs. growth chamber) in Experiment 2.

A)

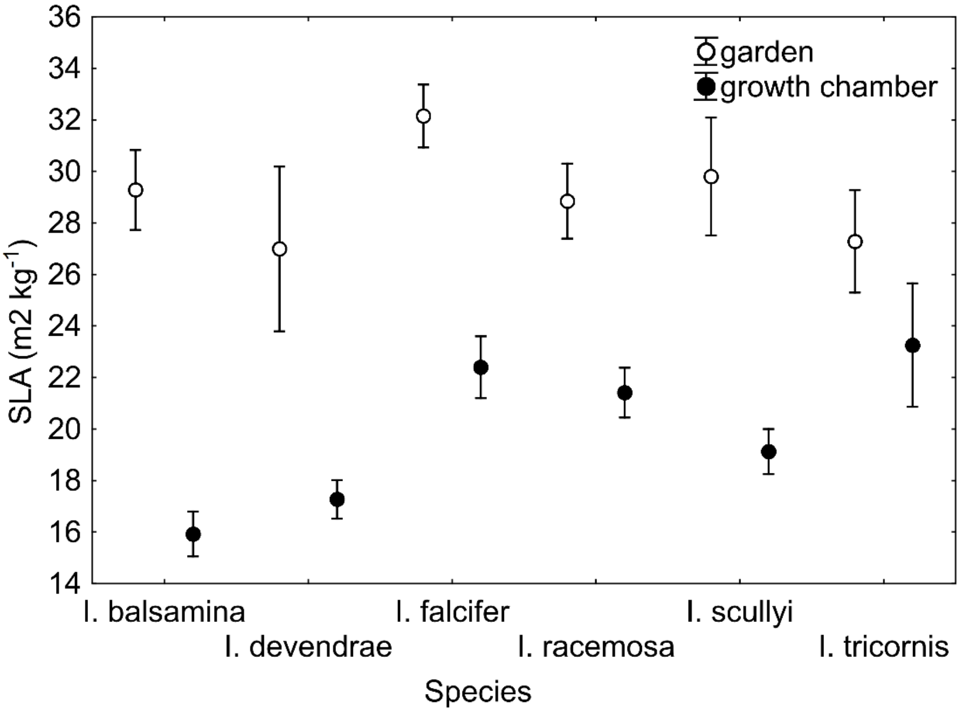


B)

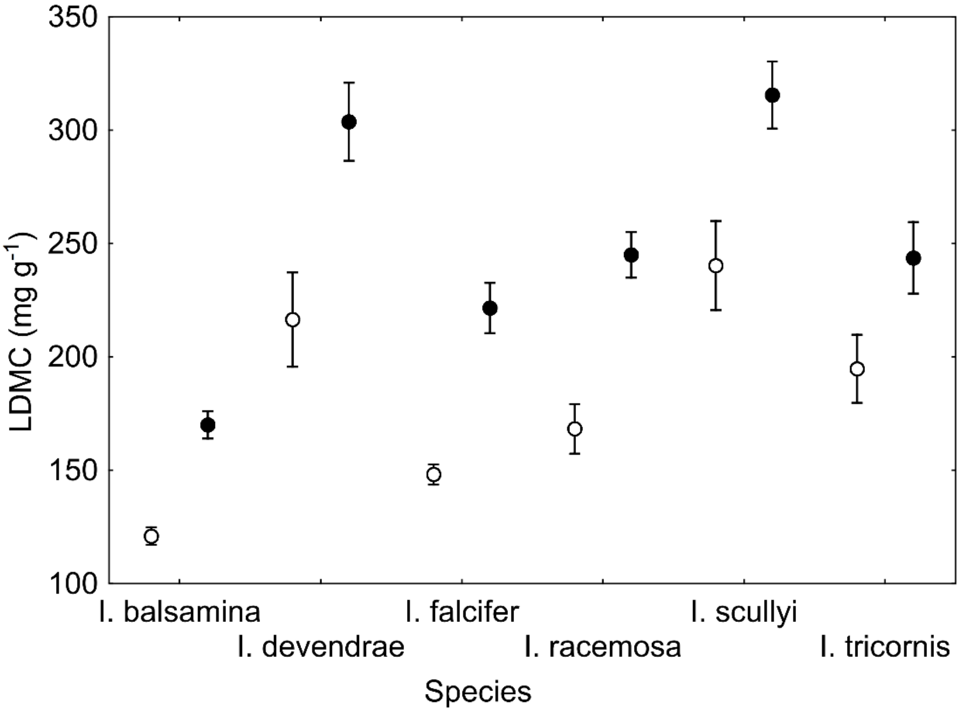

c)

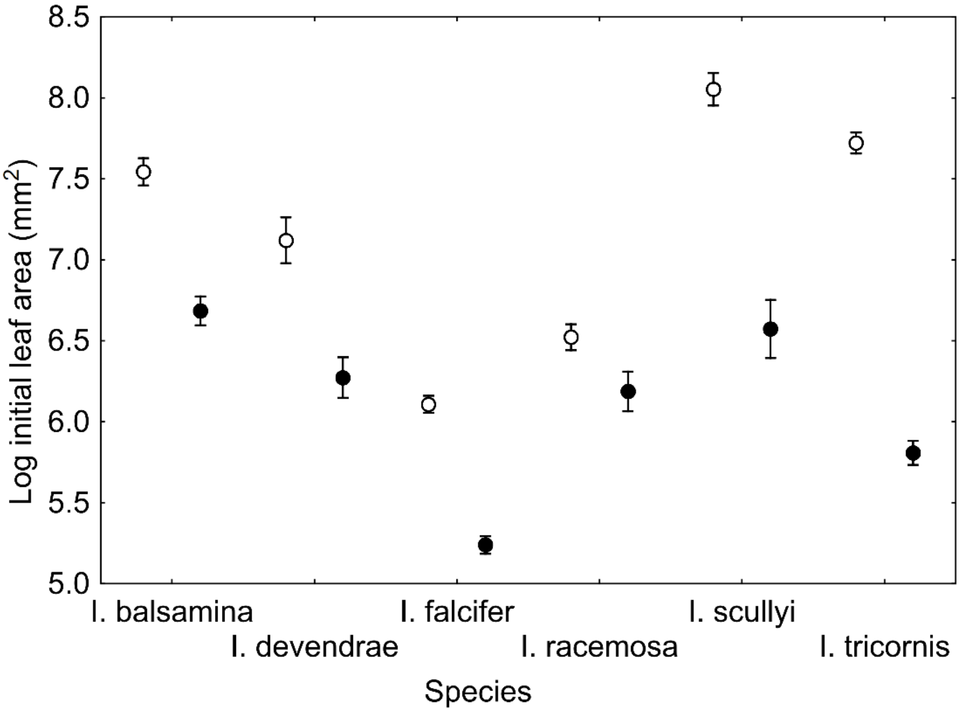
8.5
8.0
7.5
7.0
6.5
6.0
5.5
5.0
Log initial leaf area (mm2)
I. balsamina
I. devendrae
I. falcifer
I. racemosa
I. scullyi
I. tricornis
Species

**Online Resource 6** Relationship between leaf size and its nutrient content recorded at leaves both in Experiment 1 and 2.

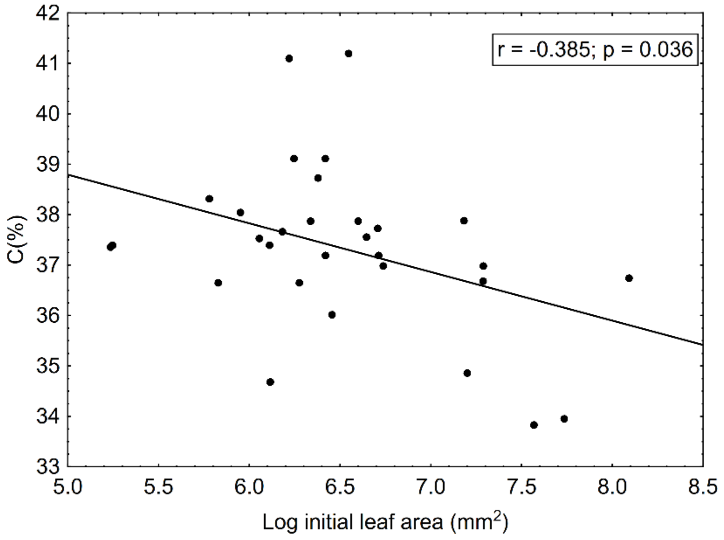


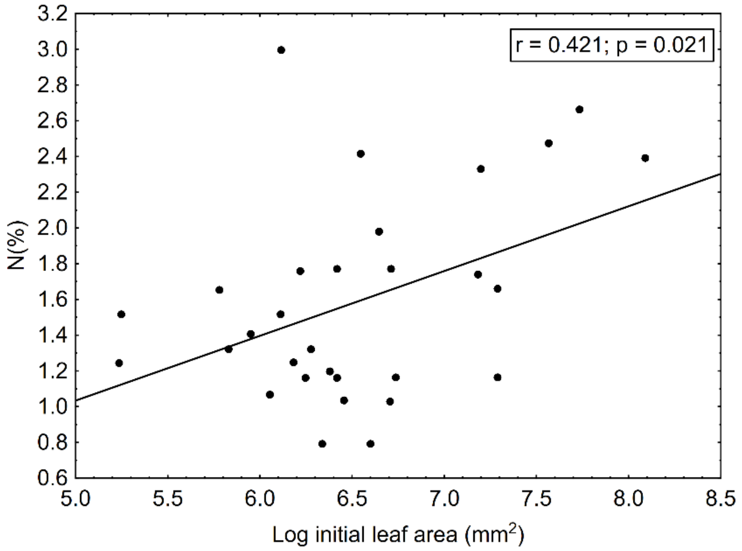


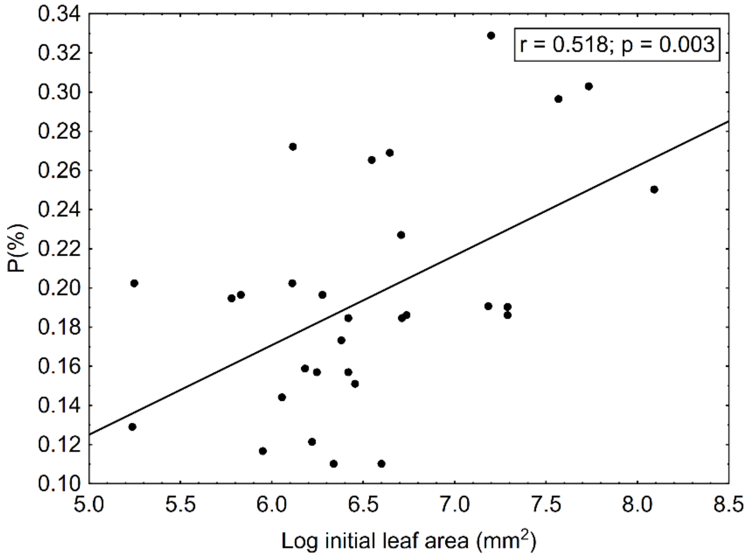
